# Genetic influences on motor learning and superperformance mutants revealed by random mutational survey of mouse locomotion

**DOI:** 10.1101/2023.06.28.546756

**Authors:** Vikram Jakkamsetti, Qian Ma, Gustavo Angulo, William Scudder, Bruce Beutler, Juan M. Pascual

**Affiliations:** Rare Brain Disorders Program, Department of Neurology; The University of Texas Southwestern Medical Center, Dallas, Texas, USA; Center for Genetics of Host Defense; The University of Texas Southwestern Medical Center, Dallas, Texas, USA; Department of Immunology; The University of Texas Southwestern Medical Center, Dallas, Texas, USA; Department of Internal Medicine; The University of Texas Southwestern Medical Center, Dallas, Texas, USA; Department of Physiology; The University of Texas Southwestern Medical Center, Dallas, Texas, USA; Department of Pediatrics; The University of Texas Southwestern Medical Center, Dallas, Texas, USA; Eugene McDermott Center for Human Growth & Development / Center for Human Genetics; The University of Texas Southwestern Medical Center, Dallas, Texas, USA

**Keywords:** Motor, CRISPR, ENU, mutagenesis

## Abstract

Evolution depends upon genetic variations that influence physiology. As defined in a genetic screen, phenotypic performance may be enhanced or degraded by such mutations. We set out to detect mutations that influence motor function, including motor learning. Thus, we tested the motor effects of 36,444 non-synonymous coding/splicing mutations induced in the germline of C57BL/6J mice with N-ethyl-N-nitrosourea by measuring changes in the performance of repetitive rotarod trials while blinded to genotype. Automated meiotic mapping was used to implicate individual mutations in causation. 32,726 mice bearing all the variant alleles were screened. This was complemented with the simultaneous testing of 1,408 normal mice for reference. 16.3% of autosomal genes were thus rendered detectably hypomorphic or nullified by mutations in homozygosity and motor tested in at least 3 mice. This approach allowed us to identify superperformance mutations in *Rif1*, *Tk1*, *Fan1* and *Mn1*. These genes are primarily related, among other less well characterized functions, to nucleic acid biology. We also associated distinct motor learning patterns with groups of functionally related genes. These functional sets included preferentially histone H3 methyltransferase activity for mice that learnt at an accelerated rate relative to the rest of mutant mice. The results allow for an estimation of the fraction of mutations that can modify a behavior influential for evolution such as locomotion. They may also enable, once the loci are further validated and the mechanisms elucidated, the harnessing of the activity of the newly identified genes to enhance motor ability or to counterbalance disability or disease.

## INTRODUCTION

No sharp boundaries can be established *a priori* between the types of functions of the organism that are primarily governed by one or a few gene products and those supported by larger ensembles of them that operate in concert. Rather, experimental evidence is required to establish, in a function-specific manner, when the characteristics of the organism are faciliated or degraded by individual genes or when they are subject to more numerous genetic influences. We set out to experimentally elucidate the relative contribution of these two possibilities to locomotion and motor learning. The motivation for the study of movement is that it represents a trainable, complex or high-order organism function where both individual genes and groups of genes may be expected to be influential. Ample precedent exists for both kinds of influence: On the reductionist end are laboratory mice subject to spontaneous or intended single-gene mutations that degrade motor phenotype. This is also the case for some human diseases that deteriorate motor ability where the causal links between gene product, cellular and organism function are few or stereotyped. In contrast, on the opposite, more complex end, most of the behaviors considered indispensable in the natural environment require adaptation to changes in initial and intervening conditions. This adaptation is often enabled by the concerted interaction of gene products that subserve the activity of the whole cell, organ or organism rather than one of their constitutent parts. The disease counterpart of this concerted action is exemplified by the dependence of mitochondrial disease manifestations on the transcriptome of a functionally related gene family rather than on individual genes (Jakkamsetti *et al*., 2022). Of note, not all individual deleterious mutations are perceptible at the level of the organism. In fact, estimation of the functional impact of damaging certain individual gene products considered critical can reveal unsuspected compensation. For example, progressive suppression or abolition of the cardiac ionic current *i*_f_, long judged fundamental to the initiation of the normal sinus rhythm and thus of the heartbeat, results in negligible disturbance of heart function due to extragenetic functional compensation by the current *i*_b, Na_ in a process entirely circumscribed to the cell membrane (Noble *et al*., 1992).

Whereas single gene discovery and characterization has been widely exploited, the investigation of concerted gene influences, which relies on the analysis of myriad genes, requires large throughput study of mutations and associated behaviors. In this context, one of the most impactful activities of the organism from an evolutive perspective is locomotion. As an eminently adaptive (i.e., subject to learning) behaviour (Jakkamsetti *et al*., 2021), locomotion is governed by an array of both well-known and poorly understood physiological and genetic influences. Locomotion is critically relevant from the perspective of human disease. For example, the motor consequences of stroke alone account for over one half of all the neurological disability in the world (World Health Organization., 2006).

Thus, we performed a forward genomic screening of mutations that may impact locomotion. Given the mutability of the laboratory mouse, we examined whether definable mouse mutations artificially induced and tested for effect in both homozygous and heterozygous states, might augment or degrade motor performance as assayed on a standard (i.e., accelerating) rotarod over several consecutive testing trials performed while blinded to genotype and compared with control mice tested simultaneously. Our work was faciliated by the fact that, originally, the C57BL/6J laboratory mouse underwent a process of stringent inbreeding, fixing alleles that might impede motor performance in the homozygous state. Indeed, many wild mice and some laboratory strains exhibit motor performance inferior or superior to C57BL/6J mice. We reasoned that most randomly induced point mutations are inconsequential, but some, by causing loss or gain of function, may alter and even augment performance at the organism level. We were particularly interested in motor superperformance mutants given the far reaching therapeutic development potential of any gain of function thus identified, since genetic gain of motor function might ultimately prove amenable to replication via the development of mechanism-related pharmacological or other interventions.

## METHODS

This study was approved by the Institutional Animal Care and Use Committee of UT Southwestern Medical Center. All other relevant institutional regulations were followed, in addition to ARRIVE (Animal Research: Reporting of In Vivo Experiments) guidelines 2.0.

### Mutagenesis

Mice were generated as previously (Wang *et al*., 2015). In brief, over approximately five years we induced germline mutations in male generation 0 (G0) C57BL/6J mice and bred them to C57BL/6J females to produce G1 (first generation after mutagenesis) males, which were subjected to whole exome sequencing to identify all non-synonymous coding and putative splicing changes induced by ENU or resident in the background. Extensive experience with ENU indicates that, with a handful of exceptions, coding/splicing changes are the main source of induced phenotype (Cordes, 2005).

While some of the mutations were homozygous lethal prior to weaning age and therefore were tested only in the heterozygous state, the majority were homozygous viable. To compute genome saturation, or the fraction of all genes destroyed or severely damaged and examined three or more times in the homozygous state, we used a method based on observation of a curated set of essential genes: those known to be essential for life based on the results of robust knockout mutations on the C57BL/6J background. We also used a larger set of ENU-induced mutations and checked the effect of mutations falling into genes of this type. This allowed us to correlate “severe damage or destruction” manifested as a lethal effect with PolyPhen-2 (Adzhubei *et al*., 2013) scores (for missense errors), or with putative null alleles (for nonsense, start lost, stop lost, critical splicing error mutations). This analysis was performed on several thousand mutations.

Lethality prior to weaning age is a phenotype with an enormous genomic footprint, encompassing large numbers of nucleotides in approximately one third of all genes. Assuming the types of damage that befall these genes are representative of the types of damage befalling all genes and the effect of that damage at the protein level, computation of the likelihood that each point mutation will cause damage can be generalized to measure the likelihood of severe damage or destruction to all individual genes, and to the sum of all genes in the genome (Wang *et al*., 2018). We concluded that, on average, only one ENU-induced mutation in six is capable of causing severe damage or outright destruction of the encoded protein. This method of estimation must be considered stringent, since certain assays are able to detect even modest damage to a protein; e.g., loss of 10% of enzymatic activity.

Average coverage of the exome target (62 million nucleotides in length) was ∼99.69% to a depth ≥10 nucleotides (quality score ≥40). Since we give consideration to two or more identical aberrant calls among 10 reads (P=0.0107 by binomial probability calculation), fewer than one in 10,000 mutations within the target sequence will be missed in sequencing. False positive mutations, while exceedingly rare, are eliminated by resequencing every non-synonymous coding/splicing change identified using Ion Torrent sequencing technology in the course of genotyping G1, G2, and G3 mice (described below).

On average, 60 coding/splicing changes were identified per G1 male. Each G1 male with ≥30 mutations and adequate breeding capacity was used to generate a large pedigree. G1 males were crossed to C57BL/6J females to yield 10 to 15 G2 daughters. Each G1 male was then crossed to its own daughters to yield ∼40 to 60 G3 mice (approximately 600-700 animals per week derived from 12-15 pedigrees), in which most of the mutations were brought to homozygosity. G1, G2, and G3 mice were genotyped at all mutation sites identified in the G1 founder, this time using Ion Torrent sequencing. In rare instances, mutations identified by Illumina sequencing were not confirmed, and were eliminated from further consideration. All validated mutations were assessed for zygosity in G2 mothers and G3 offspring. This allowed detection of probable lethal effects, evidenced by distortion of the expected Mendelian ratio. It also allowed subsequent mapping of each observed phenotype to a causative mutation, using the statistical computation method we have described (Wang *et al*., 2015).

### Gene selection for further validation

Candidate Explorer, a machine learning program developed by us (Xu *et al*., 2021), determines, across all assays of function, whether any individual mutation is likely to be validated as causative of phenotype. In brief, 60 features are used by Candidate Explorer to assess candidate mutations and score them. These features include the *p* value; the magnitude of the phenotypic effect; the number of homozygotes assayed; the variance of the data; the degree of overlap between phenotypic performance of homozygous mutant and homozygous reference allele populations; the existence of multiple alleles supporting causation; the predicted damage caused by each allele (assessed by another machine learning program); the predicted essentiality of the candidate gene (E score; assigned by still another machine learning program); and others. Candidate Explorer is trained on the basis of CRISPR/Cas9 validation data.

### Locomotor testing

We used a standard rotarod (rod diameter 3 cm, fall height 20 cm, start speed 4 rpm, acceleration rate 1 rpm/ 8 s, Touchscreen RotaRod, Harvard Apparatus) in several consecutive testing trials performed over 7 days while blinded to genotype and simultaneously testing control littermate mice. The rotarod accelerates; hence, a prolonged time to fall (or raw score) at elevated angular velocities indicates much greater motor performance than a similar prolongation at lower speeds. For example, in the case of the superpeforming mutant *Rif1^I107T/I107T^* this translates into an increase in maximum linear mouse speed of approximately from 1.0 m/ min (control) to 1.4 m/ min.

We conducted a rigor and effectiveness analysis by applying rotarod screening to mice mutated as above at a gradually increasing rate from 100 to 900 mice per week by a team of six testers who tested an approximately equal number of mice per week. Testing more than 900 mice per week nearly exceeded testing capacity and was thus considered prohibitive for efficient screening. Conversely, low mouse numbers also negatively impacted phenotypic mutation detection rate. **Supplementary figure 1** illustrates the effect of varying the number of mice per week relative to the number of genes with likely deleterious (probably damaging or null) mutations studied each week. Increasing weekly mouse numbers screened exerts a sharp impact on the number of mutant genes studied for each particular week. Thus, we estimated the optimal rate of screening at about 600 (but fewer than 900) mice per week, which was ultimately contingent upon weekly mouse generation rates (subject to unavoidable biologically-imposed fluctuations) and can be adjusted to accommodate pedigrees that span about 100 more or fewer mice.

To make the behavior comparable across testers and time, we converted raw rotarod fall latency scores to normalized scores for gene linkage analysis and T-scores for subsequent motor behavior analysis. Normalized scores were obtained by first deriving a scaling factor by dividing 100 by the mean wild type mouse performance in that week of testing (**supplementary figure 2**). The scaling factor was multiplied by the raw score of each mouse (in wild type, REF (reference non mutated mice littermate to ENU mutagenized mice), HET (heterozygous mutants) and VAR (mutant homozygous) groups) tested that week to calculate their respective normalized score. The mean normalized wild type mouse score was thus 100 for each week and the REF, HET and VAR mice scores in relation the wild type mean could be compared across weeks. T-scores (calculated as 50+(10×Z-score)) were derived from Z-scores (which are often used for similar forward genetics projects (Kumar *et al*., 2011; Cinà *et al*., 2019)) but modified to allow mathematical operations on non-negative numbers for standardizing data across time and locations (Levin *et al*., 2020). T-scores also allowed for a simple identification procedure of outlier performance since the mean is set at 50 and every 10 units correspond to one standard deviation. For control, untrained wild type mice were tested each week alongside the ENU mutagenized mice totaling 1,408 mice. Rotarod T-scores for the 6^th^ trial in these mice tested over time showed stability (**supplementary figure 2**). As expected for wild type mice, most of the performances were within 2 standard deviations of the mean and a few were within 3 standard deviations.

### Principal component analysis (PCA) and gene enrichment analysis

PCA was conducted on the motor behavior parameters listed in **supplementary table 1**. Overall, the parameters reflected rotarod performance scores at each trial, learning between trials and likely learning trajectories including early and late rise in performance and learning rate. To allow comparison across multiple continuous parameters of varying magnitude, the values for each parameter were normalized to range from 0 to 1 prior to conducting PCA. Gene enrichment analysis (GEA) was conducted using EnrichR (Kuleshov *et al*., 2016). To examine initial learning curves over just 6 trials, T-score values were smoothed with a median filter over 2 adjacent trials. Statistical analyses examining differences between groups employed original T-scores prior to smoothing.

## RESULTS

### Genome coverage (saturation) rate

A total of 36,444 such mutations were screened (in either heterozygous or homozygous state). 32,075 of these mutations resided in a total of 32,726 G3 mice, which were screened in the homozygous state at least once. By computing gene damage or destruction, we estimate that 16.3% of all autosomal genes were severely damaged or destroyed by mutations that were brought to homozygosity in five or more G3 mice, each tested six times using the rotarod. Note that the X chromosome is not mutagenized in our G3 mice: it was derived from a wild type C57BL/6J female mated to a mutagenized G0 male, which contributed a Y chromosome to the G1 founder. However, spontaneous mutations of the X chromosome did occur and were detected. If all chromosomes are included in the saturation estimate, at least 11.4% of all genes were damaged or destroyed by mutations that were brought to homozygosity in five or more G3 mice, each tested six times using the rotarod. The average percent of each mutation effect was: Probably benign: 38.3%, possibly damaging: 15.1%, probably damaging: 38.8%, and probably null: 4.5%. The remaining 3.3% can be accounted for by mutations classified as “unknown”.

### Superperformance genes

By the criterion of “good or excellent” candidate ranking in Candidate Explorer (Xu *et al*., 2021), we noted 15 mutations in 14 genes that either increase or decrease rotarod performance (**table 1**).

**Table 1.**
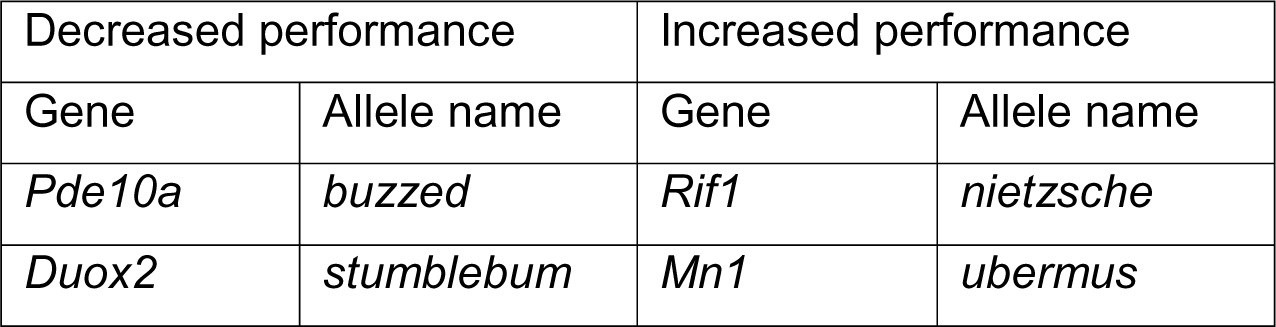
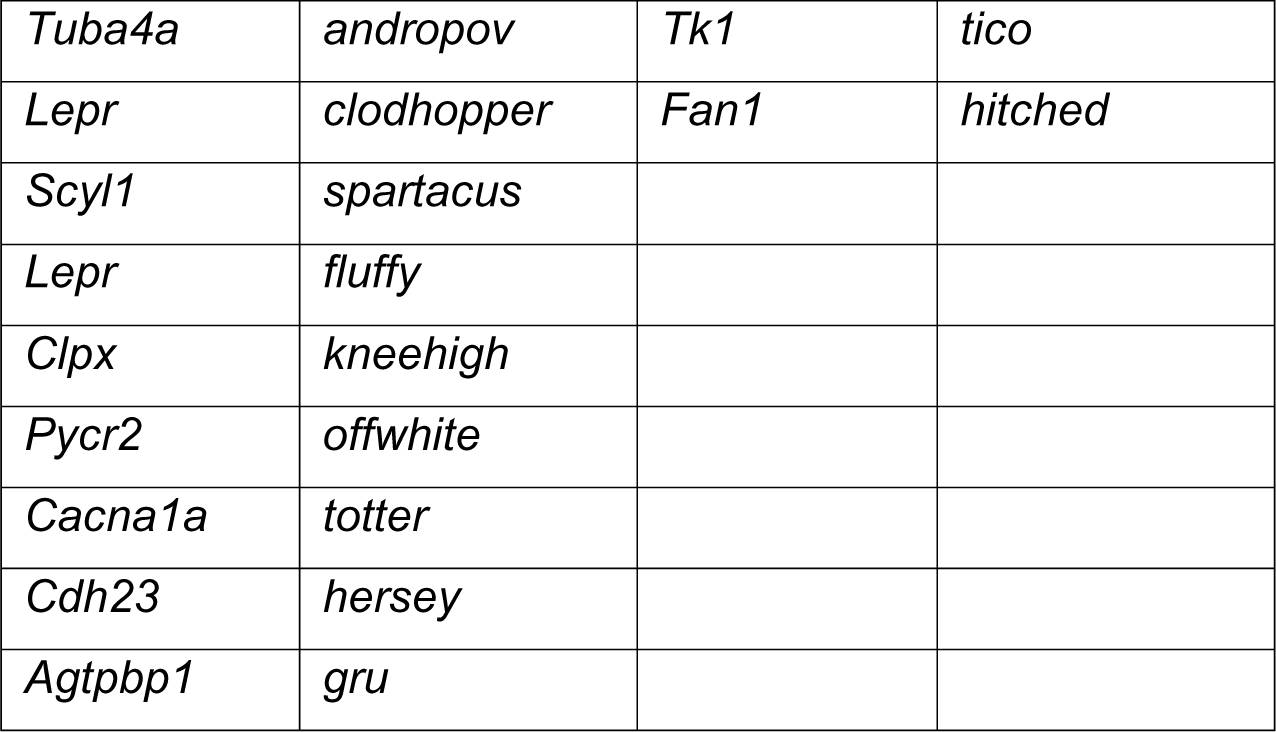
Sub and superperforming alleles uncovered by mutant mouse rotarod motor testing.

Several of these loci are known causes of brain diseases with ataxia or dystonia. All are recessive except for *struggles*, an allele of *Nfatc3* that is also associated with reduced survival. All are unambiguously assigned to a single mutation except *even-steven*, a phenotype that results from one of two co-segregating mutations, in *Ak8* or *Ntng2* (Candidate Explorer favored the former). Two separate mutations in the leptin receptor (*Lepr*) gene, both leading to obesity, caused poor rotarod performance. Although *Lepr* is expressed in the brain, obesity mutations (including *Lep*, which is not brain-expressed) can cause poor rotarod performance. Among the mutant genes that enhance rotarod performance, we unambiguously assigned (Candidate Explorer verification probability, CEvp= 0.974) a single recessive ENU-induced mutation, *Rif1^I107T/I107T^* (*nietzsche*) that increases rotarod performance (**figure 1A**). We also discovered *Tk1^V140E/V140E^* (CEvp= 0.689, *tico*), *Fan1^I618N/I618N^* (CEvp= 0.698, *hitched*) and *Mn1* (CEvp= 0.796, *ubermus*) mutations in relation to motor superperformance (**figure 1B-D**).

**Figure 1.**
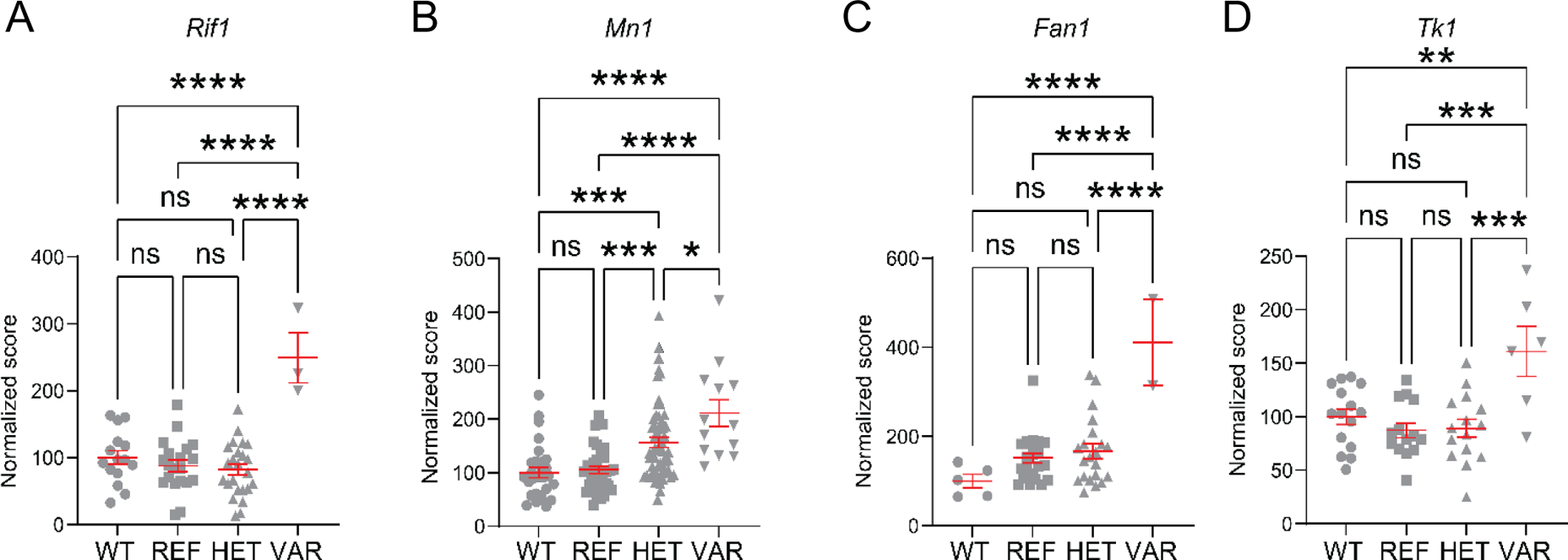
Rotarod normalized scores for superperforming mice. Specific mutations in genes *Rif1* (A), *Mn1* (B), *Fan1*(C) and *Tk1*(D) induce a motor phenotype of superior performance on the rotarod test. The scores correspond to the 4th trial for *Mn1* and *Tk1* and the 6th trial for *Rif1* and *Fan1*. Abbreviations: WT: wild type, REF: reference non mutated mice littermate to ENU mutagenized mice, HET: heterozygous mutants, VAR: variants with homozygous mutations. n.s. non significant comparison; *: p< 0.05; **: p< 0.01; ***: p< 0.005; ****: p< 0.001.

### Functional role of four superperformance genes

We examined brain microarray data from the Allen Brain Atlas (Lein *et al*., 2007) for the regional expresion of the superpeformance genes (**figure 2**). *Fan1* displayed greater expression in pons or medulla, *Mn1* in striatum, *Rif1* in cerebellum and *Tk1* in the medulla. Overall, these structures are generally relevant to motor performance. There was a distinct functional role common to these 4 genes. Gene enrichment analysis indicated a significant association of these genes with nuclei acid biology and DNA repair and modulation. Gene Ontology Molecular Function indicated that this gene set’s association with nucleoside kinase activity (**figure 2B**) was higher than that of randomly selected genes.

**Figure 2.**
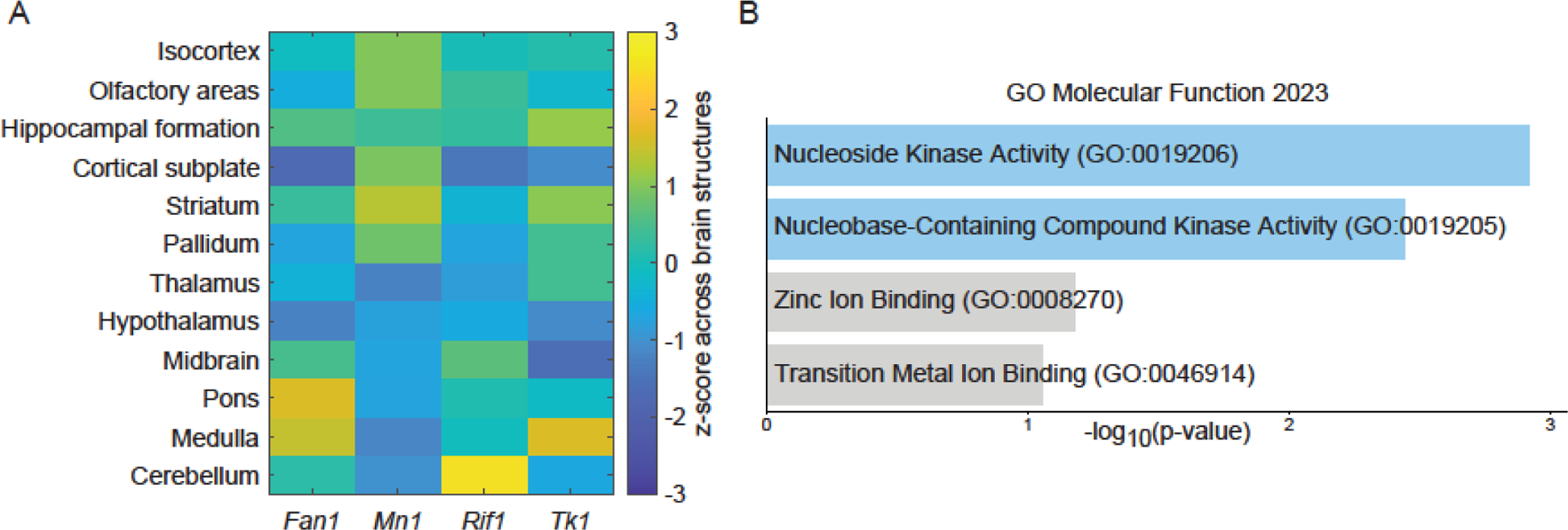
Brain expression and overall function of 4 superperformance genes. A. Allen Brain Atlas’ *in situ* hybridization summary data for brain structures in the mouse. B. Gene enrichment analysis of function. Blue shadowing indicates significance (p< 0.05) for adjusted p-values.

### Distinct clusters of learning and non-learning ENU mutagenized mice

We also examined the impact of ENU mutations on motor learning. The first few trials during rotarod training involves rapid learning for the mouse, associated with learning to regulate foot position and speed to adapt to the accelerating surface (Buitrago *et al*., 2004; Shiotsuki *et al*., 2010; Jakkamsetti *et al*., 2021). Evidence of such learning is manifest as early as after a single rotarod trial (Jakkamsetti *et al*., 2021). When compared to wild type, the mice with ENU mutations performed slightly worse than wild type (**figure 3A**). This is in agreement with previous studies (Eyre-Walker & Keightley, 2007) that indicate that the majority of mutations can have a neutral or deleterious impact with very few providing an advantage.

**Figure 3.**
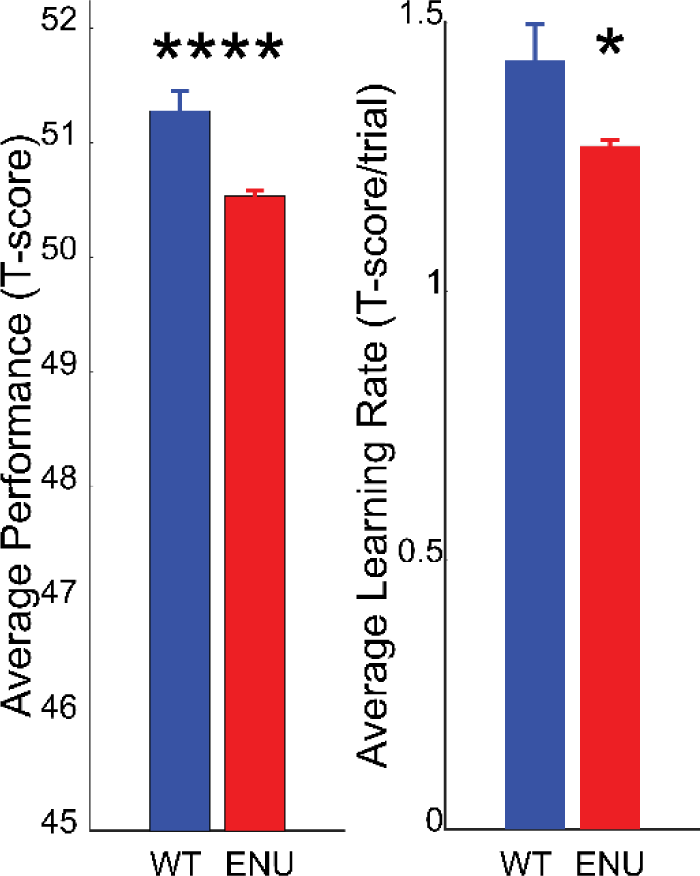
Average rotarod performance (left) and average learning rate (right) for wild type and ENU mutagenized mice.

Average motor learning rate was also reduced for ENU mice (**figure 3B**). However, considering the large number of ENU and wild type mice tested, it was surprising that the difference in their respective learning rates barely reached statistical significance (p= 0.026). This suggested that there was considerable variability in the average learning rate and that a significant number of mice exhibited values overlapping with wild type, thus decreasing statistical significance. Hence, to test whether behavioral clusters corresponding to different learning patterns across 6 trials existed, we conducted an unsupervised principal component analysis clustering of data with multiple parameters including those that reflected different learning trajectories (**supplementary table 1**). This revealed two main clusters of mouse performance, one comprising 20,672 mice and one with 1,837 mice (**figure 4A**). Compared to the smaller cluster, mice from the larger cluster displayed learning trajectories similar to wild type mice (**figure 4B**). In contrast, mice from the smaller cluster displayed flat learning curve trajectories. The smaller cluster mice did not exhnibit a net positive learning rate and, in fact, were slightly below a learning rate of zero (mean= −0.11, p< 0.0001, one sample t test against zero) and were significantly different from wild type mice and mice from the larger cluster in their average learning rate (**figure 4C**). Thus we henceforth refer to the larger cluster as the learning group mice and the smaller cluster as the non-learning group mice. The first nascent baseline performance for the smaller cluster was higher (**supplementary table 2**), and the final performance score at the 6th trial was lower than that of mice in the larger cluster. We noted that the average performance over 6 trials was slightly lower than wild type for both ENU mutagenized learning groups, and more so for the non-learning group (**figure 4D**).

**Figure 4.**
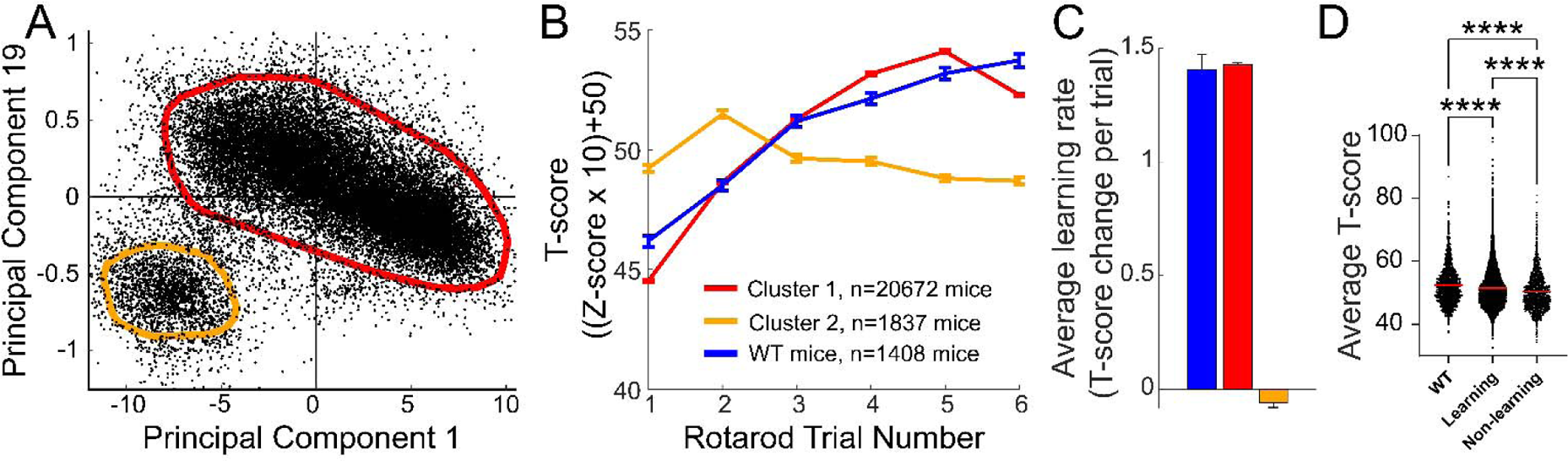
Motor behavioral cluster of ENU mutagenized mice. A. PCA analysis of mouse rotarod behavior reveals two distinct clusters. The thick red and orange lines were determined by density-based clustering algorigthm DBSCAN. B. Rotarod performance across 6 trials for the two clusters isolated in (A) and wild type (WT) mice. C. Average learning rate across 6 trials for the two clusters isolated in (A) and wild-type mice. The color of bars represent the same groups as the colors in (B). D. Average performance across 6 trials.

### Mutation rates in learning and non-learning mouse groups

Since motor learning is, arguably, critical for survival, we hypothesized that the mice that did not learn would harbor a higher number of homozygous mutations given that, in general, mutations have no impact or cause deficits in behavioral performance. Surprisingly, the the non-learning mice had fewer mutation numbers compared to the learning group (**figure 5A**). This prompted us to re-examine our hypothesis that more mutations would result in deteriorating motor performance. Thus we examined the impact of the number of mutations in a mouse on performance. However, it is possible that, with each cumulative mutation in a single mouse, the performance will eventually begin deteriorating in some animals at some mutation number, thus confounding the analysis. Therefore, we first examined the coefficient of variation of performance across mutation rates and selected 15 as the number of mutations beyond which performance variability changed drastically (**supplementary figure 3A**). We found that, as expected, total motor performance deteriorated slightly but significantly as the number of mutations increased (**supplementary figure 3B**), yet the relationship between the two was weak and the dot plot of average t-score performances against mutation number did not show any significant relationship (**figure 5B**). Interestingly, the initial performance of the mouse (i.e., the average of the first two trials) was more correlated with deterioration upon increase in mutation number. Similarly, the average learning rate increased slightly but reliably with increase in number of mutations in a mouse (**figure 5C, supplementary figure 3C**). There was an opposite relationship between initial motor performance and learning rate as expected. Mice that started with low initial performances demonstrated better learning (**figure 5E**). The 6,096 genes with homozygous mutations that were unique to the learning group (**supplementary table 3**) shared a common role in Gene Ontology Molecular Function 2023 that indicated DNA modulation with histone methyltransferase activity (**figure 5F**).

**Figure 5.**
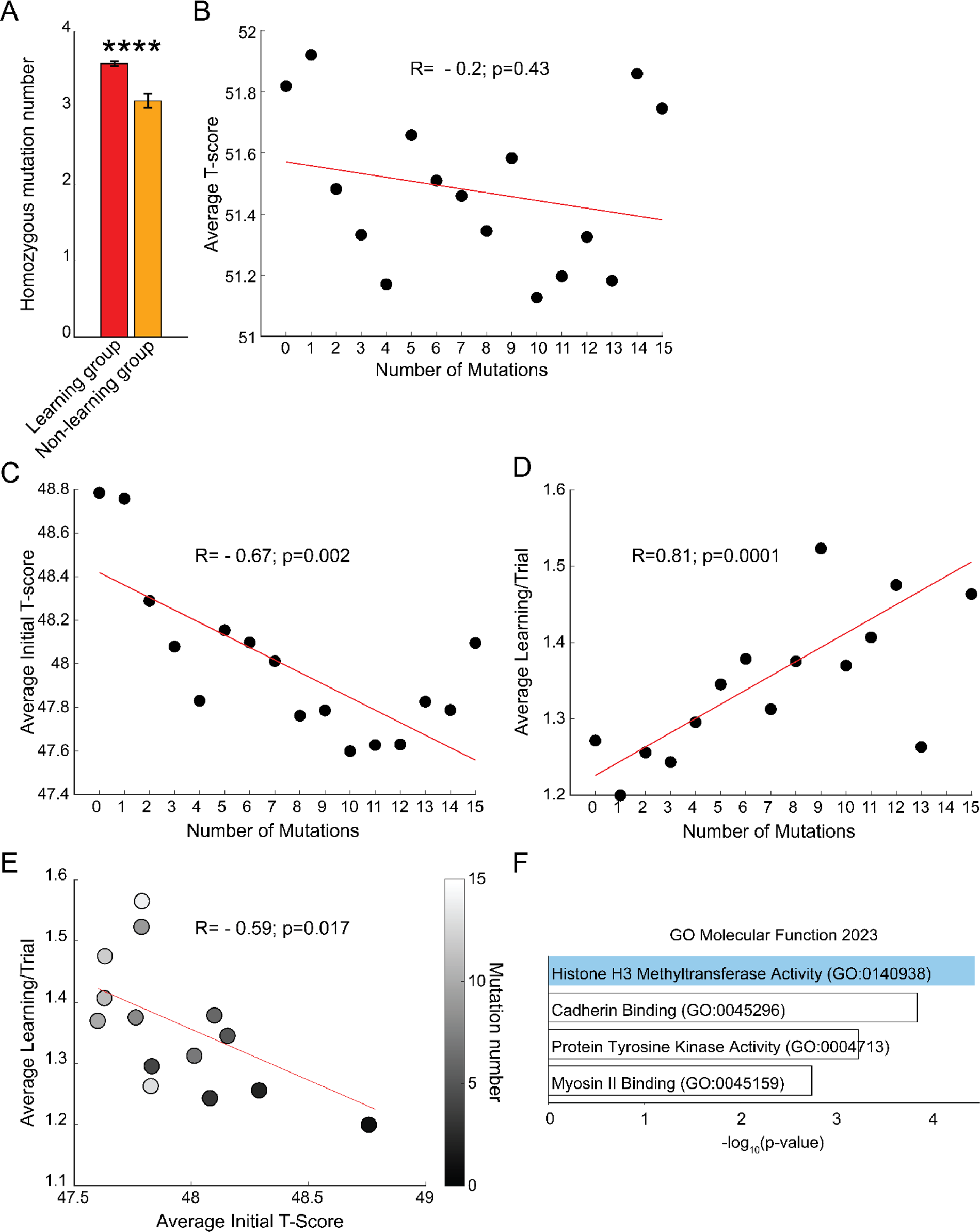
Mutation rate for behavioral clusters. A. Homozygous mutation number per mouse in learning and non-learning mouse groups. B. Mean of average t-scores for a given number of mutations in a mouse (a dot plot for all mice is in **supplementary figure 3**). C. Mean of initial t-scores for a given number of mutations in a mouse. D. Mean of average t-scores for a given number of mutations in a mouse. E. Mean of initial t-scores for a given number of mutations in a mouse plotted against corresponding mean learning rate for the same mutation number. F. Gene enrichment analysis of mutated genes unique to the learning group. Blue shadow indicates significance (p< 0.05) for adjusted p-values.

### Number of mutations required for for superperformance

The combination of several mutations can confer a behavioral advantage. Therefore, we estimated the mutation rate that could enable the emergence of a superior or outlier motor behavior. Based on the wild type mouse performance data where no mice deviated more than 3 standard deviations from the mean, we defined outlier performance as an increase in performance greater than 3 standard deviations. We found 39 mice with outlier performance, amounting to 0.15% of mice (**figure 6A**). In other words, for approximately every 672 ENU mice there was an outlier performing mouse. Given the total number of homozygous mutations harbored by ENU mice, one mouse became an outlier in motor performance on the rotarod approximately for every 2,410 homozygous mutations. Regarding the learning rate we fould 696 mice with outlier performance (**figure 6B**) amounting to 0.27 % of mice. In other words, for every 38 ENU mice one was a motor learning outlie, or one mouse became an outlier in motor learning rate per approximately 696 homozygous mutations studied.

**Figure 6.**
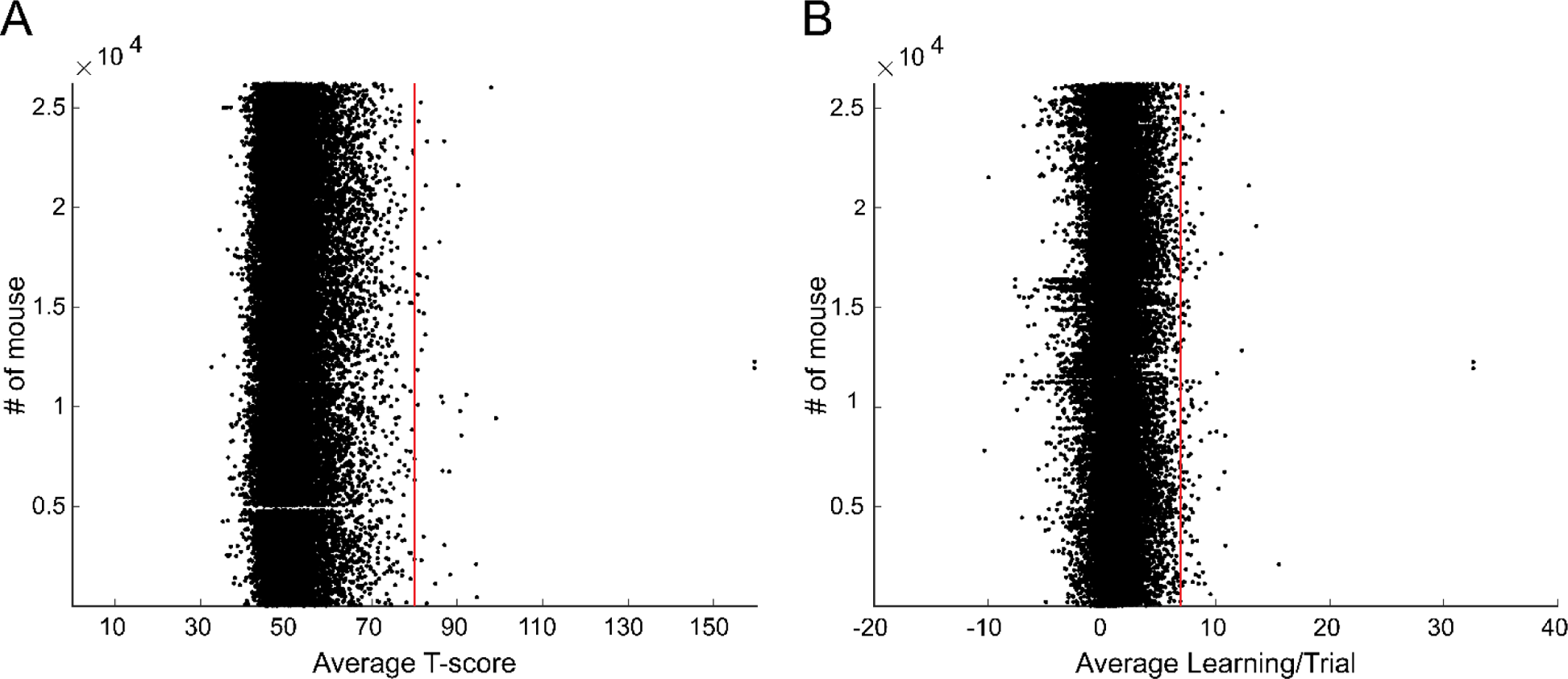
Mutation rate associated with outlier motor performance. A. Average T-score for each mouse stacked longitudinally. B. Average learning for each mouse stacked longitudinally. Red lines demarcate 3 standard deviations above the mean.

### Relationship between number of mutations and survival

Elevated mutation rates can be expected to reduce survival. Thus, we examined the relationship between mutation number and post-weaning survivability. There was a significant logarithmic relationship between mutation load in a mouse and survivability (**supplementary figure 4**), with the number of surviving mice decreasing logaritmically as the number of mutations in a mouse increased.

## DISCUSSION

### Scale and effort-yield ratio relative to alternative approaches

While in principle individual knockout mutations could be generated one at a time to cover the genome and tested to identify those that enhance motor performance, this could not possibly be accomplished as efficiently as a screen of ENU-induced mutations: First, assuming there are ∼25,000 genes in a mouse (Guenet, 2005), targeted testing of each gene (using, for example, 20 mice per mutant pedigree) would require 500,000 mutant mice, which would take over 16 years if 500 mice per week are tested. Second, such an effort would not permit a comprehensive assessment of all genes: About 1/3 of all genes are essential for survival to weaning age, and ENU permits analysis of this fraction of the genome by inducing viable hypomorphic alleles, which regularly create living mice with measurable phenotypes. Moreover, in each pedigree, an average of 60 ENU-induced coding/splicing changes are examined in all zygosities. The causative mutation is usually identifiable using the automated mapping software of Mutagenetix (Linkage Analyzer), given a panel of 30-50 G3 mice. Each week 500 to 700 mutations can be examined in this way, within a total of 7-10 pedigrees. The alternative, i.e., the expansion of 600-900 single pedigrees per week, each bearing a mutation and tested for our purposes, would be impractical. Even if only 20 mice were generated per pedigree, we would not be able to phenotype 12,000 to 18,000 mice per week. The multiplex analysis permitted by the ENU approach makes it much more powerful than the single mutation approach.

The efficiency of our method (15 super- or sub-performance motor genes identified after ∼30,000 mice studied - or one gene for each ∼2000 mice) is comparable to other reports that also use ENU mutagenesis and behavioral screening for sleep (Funato *et al*., 2016) (2 behavior-modifying gene mutations reported for ∼8000 mice) or by comparing mice of different strains (Kumar *et al*., 2013) (1 gene mutation reported for ∼1000 mice).

In summary, whereas several years of effort were once needed to determine the source of a phenotype via positional cloning, we have made it instantaneous. When a phenotype is detected in screening, its cause is generally known at the same time. We have also greatly increased the sensitivity with which quantitative trait mapping can be performed. Whereas formerly only phenotypes with a large effect size could reliably be ascribed to a causative mutation, we show that it is now feasible to instantly map mutations with effect sizes of a few percent, in which there is considerable overlap between the phenotypic performance of wild type, hetero and homozygous populations. These conditions have transformed forward genetics and made it possible to reliably determine which mutations cause increases as well as decreases in rotarod scores. This type of work cannot be accomplished in human populations, in outbred mice, or in recombinant inbred mapping projects, where mutation density greatly exceeds the density of ENU-induced mutations.

### What is to be considered ‘superperformance’?

As with the term ‘gain of function’, ‘superperformance’ depends on the type of ‘performance’ that one refers to, since this term applies to various biological levels of observation. For example, increased time on a rotarod may be due to greater paw movement velocity, or decreased variance of paw position at low rod speed, or even to the opposite variance at high speed. In several publications (Piochon *et al*., 2014; Vergouts *et al*., 2015; Sheppard *et al*., 2022), the relationship of these parameters with, for example, linear gait velocity may not be straightforward or even appear counterintuitive. For a mouse with a broad-based gait will do well on a slippery rock but poorly on a narrow liana. Thus, the quantification of motor performance upon a complex motor behavior such as rotarod rate of learning and time to fall is likely to prove more translatable to humans than a simpler task (Jakkamsetti *et al*., 2021).

### Function of the genes associated with enhanced motor performance or motor learning

We identified *Rif1*, *Tk1*, *Fan1* and *Mn1* as genes that, when mutated, lead to motor superperformance. We have also uncovered an additional 6,096 genes associated with a faster than expected (i.e., relative to other mutant mice) motor leaning rate.

The available information about the normal function of *Rif1*, *Tk1*, *Fan1* or *Mn1* is uneven. For example, whereas these genes are differentially expressed in brain areas potentially relevant to locomotion and whereas all have been implicated in nucleoside phosphorylation, more significant evidence is available regarding the function of *Rif1*. *Rif1* regulates several DNA damage response processes, particularly in relation to double stranded break (DSB) repair, as it influences repair pathway selection between non-homologous end-joining (NHEJ) versus homologous recombination repair (HRR) (Scully *et al*., 2019). DNA repair is related to synaptic strength: Loss of polymerase µ function can induce less efficient but more conservative NHEJ repair, resulting in increased mitochondrial respiration efficiency and enhanced hippocampal long term potentiation (Lucas *et al*., 2013), which may underlie synaptic potentiation evoked by repeated motor activity (Whitlock *et al*., 2006), whereas DSB formation may be a physiological event that leads to early-response gene expression relevant for experience-driven changes to synapses (Madabhushi *et al*., 2015). Additionally, RIF1 influences chromatin function by directly binding to G4 DNA structures (Kanoh *et al*., 2015). *Rif1* regulates the formation of telomeric and transcriptional RNA-DNA hybrid R-loop structures, which are sources of endogenous DSB (Tubbs & Nussenzweig, 2017) relevant to human diseases including neurological ones (Richard & Manley, 2017). Thus, the *Rif1* mutation might potentiate DNA repair and/or alleviate inhibitory G4 architecture leading to unknown changes in transcript expression that result in enhanced motor peformance.

A salient association of the 6,096 genes related, when mutated, to relative accelerated motor leaning is with histone H3 methyltransferase activity. Whereas numerous additional activities may be relevant given the large number of these genes, mouse histone regulation via methylation has been implicated in certain forms of synaptic plasticity (Shen *et al*., 2016), including hippocampal changes in relation to habituation to novel environments (Collins *et al*., 2019).

In summary, whereas we might have entertained *a priori* that ion channel and other membrane function genes would be best situated to exert the changes of neuronal function necessary for superperformance or increased motor learning, we find that the most robust associations point to DNA regulatory aspects.

### Evolutionary implications

Given the rate of locomotor-impacting mutations we have identified and the estimated spontaneous mutation rate, it is possible to conjecture the probability of appearance of a modification in locomotor behavior over time. **Table 2** illustrates the estimated spontaneous mutation rates of several organisms. The table reflects wide variation and the values should be taken with further caution because the genetic background likely influences the mutation spectrum, in addition to a potential for segregating variation among inbred mouse strains (Dumont, 2019). If we take one estimate from **table 2**, the probability of a homozygous mutation happening by chance through spontaneous mutations is (7.9×10^−9^) × (7.9×10^−9^) or 6.24×10^−17^. Taking into account our results indicating that 0.15% and 0.27% of all the homozygous ENU mutations studied modify locomotor performance and motor learning phenotype, about (7.9×10^−9^) × (7.9×10^−9^)×0.15) mutations, or 9.36×10^−18^ (1.19×10^−7^%) of mutations per replication may endow a superior phenotypic impact. Similarly, 2.13×10^−7^ % of mutations per replication may endow superior motor learning.

**Table 2.**
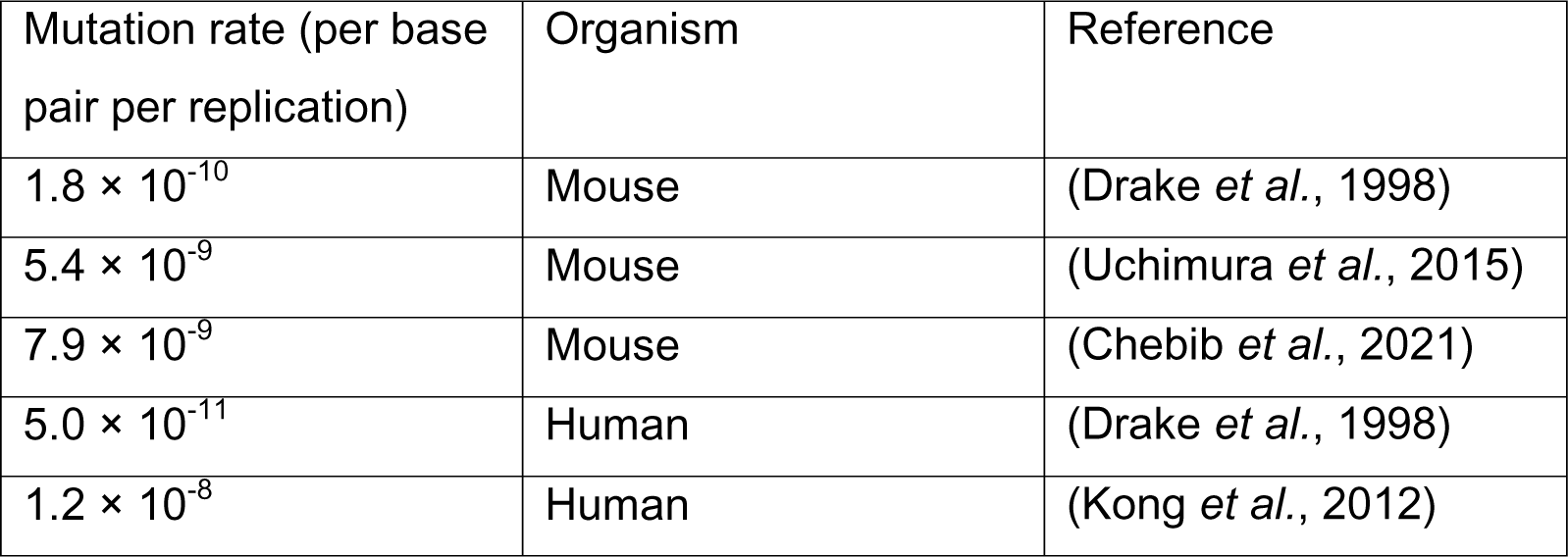
Spontaneous mutation rate estimates in human and mouse organisms.

| Mutation rate (per base pair per replication) | Organism | Reference |
| --- | --- | --- |
| $1.8 \times 10^{-10}$ | Mouse | (Drake *et al.*, 1998) |
| $5.4 \times 10^{-9}$ | Mouse | (Uchimura *et al.*, 2015) |
| $7.9 \times 10^{-9}$ | Mouse | (Chebib *et al.*, 2021) |
| $5.0 \times 10^{-11}$ | Human | (Drake *et al.*, 1998) |
| $1.2 \times 10^{-8}$ | Human | (Kong *et al.*, 2012) |

### Conclusions

Despite the ever-increasing diversity of neurological disease mechanisms elucidated, motor impairment remains a common and poorly treatable outcome. For example, the motor consequences of stroke alone account for over one half of all the neurological disability in the world (World Health Organization., 2006). In contrast with this, some persons and animals are endowed with supernormal motor ability stemming from spontaneous mutations in genes such as myostatin (Schuelke *et al*., 2004). Disorders as different as muscular dystrophy and spinal muscular atrophy benefit from myostatin blockade in mice (Bogdanovich *et al*., 2002; Long *et al*., 2019), suggesting that genetic or pharmacological modulation outside the primary disease locus can mitigate motor impairment. Our results suggest that superperformance can be identified at a measurable rate. After further mechanistic work, it may be possible to induce it to augment limitations in motor peformance or recovery, thus mitigating illness or disability.

## AUTHORS’ TRANSLATIONAL PERSPECTIVE

The discovery of useful *drugs that augment motor function* is not new. For example, physostigmine, a reversible cholinesterase inhibitor, treats myasthenia gravis by increasing the amount of acetylcholine available at the neuromuscular junction. The development of physostigmine as a drug was built upon studies of the actions of a paralyzing poison from the Calabar plant (*Physostigma venenosum*) over the course of six decades. Rather than directly searching for new chemical matter with beneficial pharmacological effects, we conducted a genetic search in the laboratory mouse for those proteins in which mutation (generally leading to loss of function similar to that caused by a drug), leads to enhanced motor performance.

## Data availability statement

All the mouse screening data are freely available from Mutagenetix (https://mutagenetix.utsouthwestern.edu) and the mice from the Mutant Mouse Resource & Research Centers (https://www.mmrrc.org/). The additional *Rif1* mutant behavioral data presented are available from JMP.

## Competing interests

None

## Author contributions

Conception and design: JMP, BB

Acquisition, analysis, and interpretation of data: All authors

Drafting of the manuscript: JMP, BB, VJ

Critical revision of the manuscript for important intellectual content: All authors

All authors approved the final version of the manuscript, agree to be accountable for all aspects of the work regarding the accuracy or integrity of any part of the work and qualify for authorship. All of those who qualify for authorship are listed.

## Funding

National Institute of Neurological Disorders and Stroke grant NS121993 (JMP, BB)

### Acknowledgments

We thank Sohini Mukherjee, Nanda Regmi, Lauren Bailey, Doretha Evans and Katrina Young for conducting mouse motor tests.

## Supplementary information

**Supplementary figure 1.**
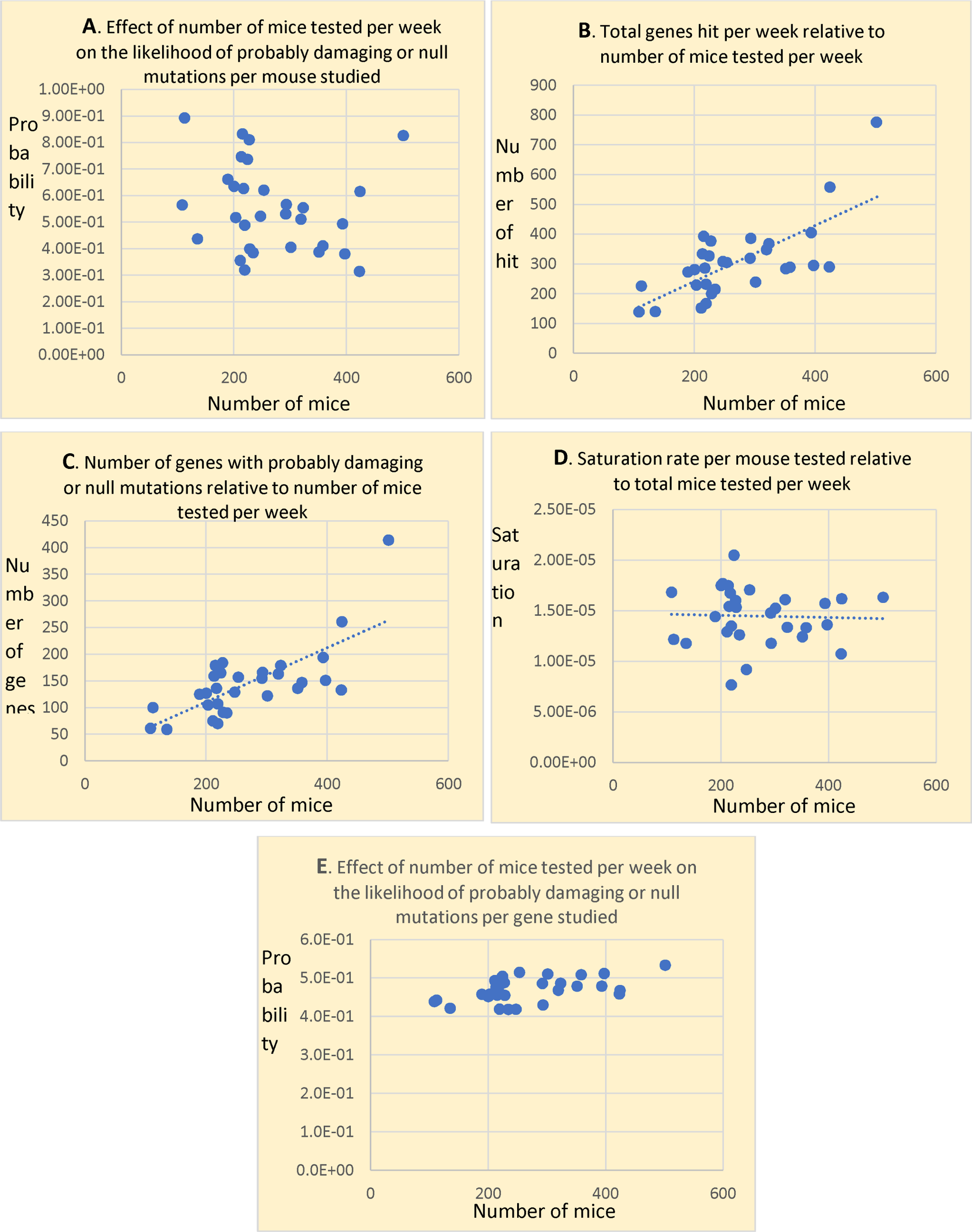
Efficiency of the motor screening. The genes affected include probably benign, possibly damaging, probably damaging and probably null mutations.

**Supplementary figure 2.**
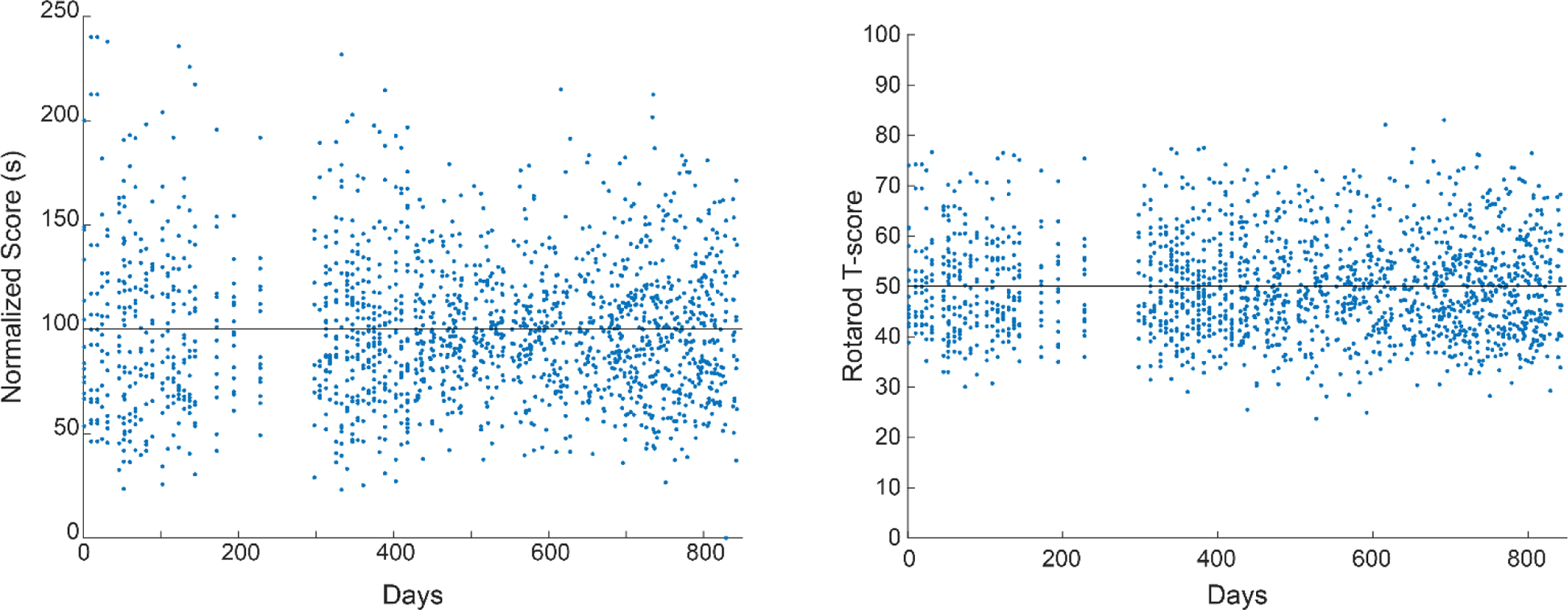
Normalized rotarod scores (left panel) and T-scores (right panel) of control wild type mice over time. Most rotarod T-scores lie within mean +/− two standard deviations (50 +/− 20) and almost all T-scores lie within 3 standard deviations of the mean (50 +/− 30).

**Supplementary figure 3.**
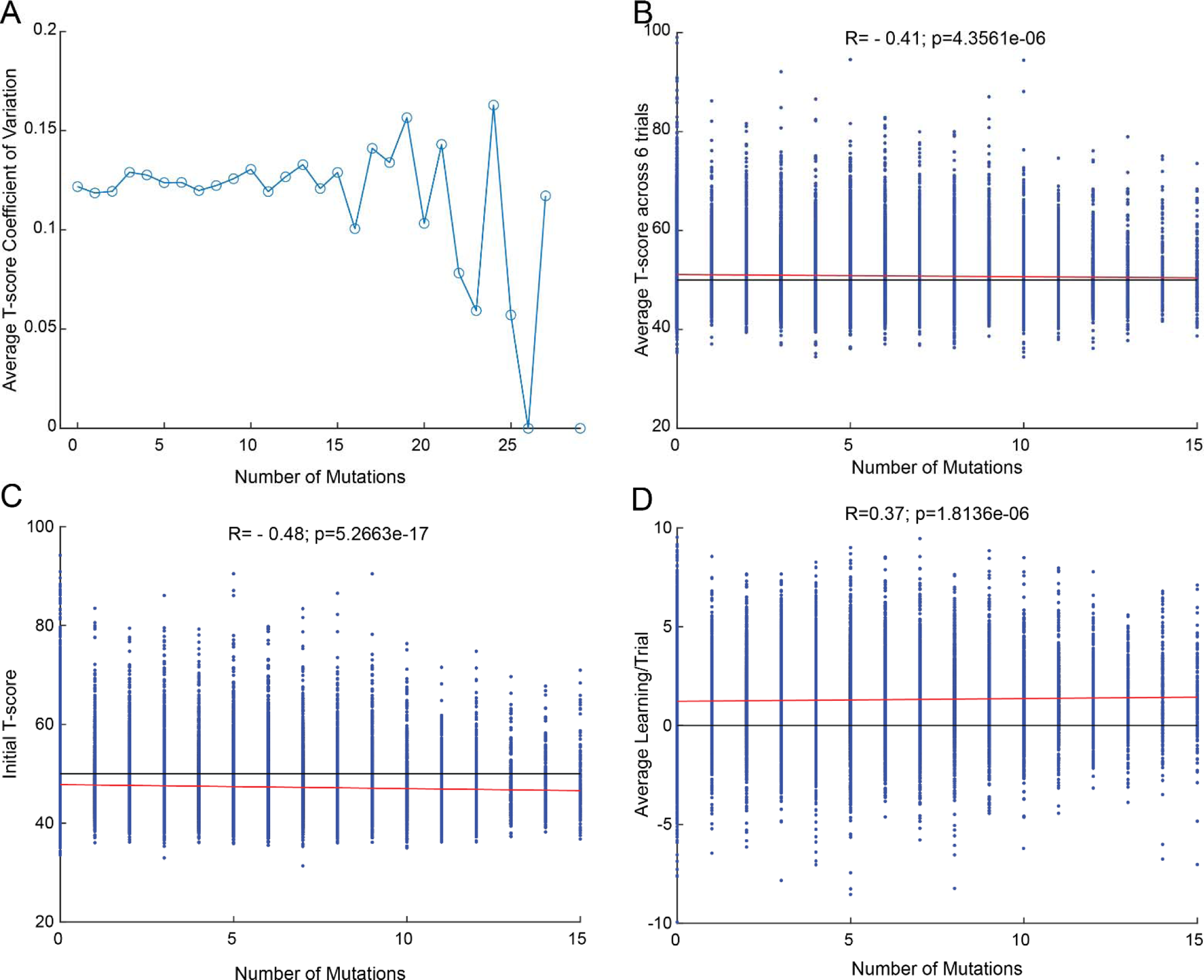
Impact of mutation rate on rotarod performance. A. Coefficient of variation of T-scores for a given number of homozygous mutations in a mouse. Note a relatively large negative change in variation after 15 mutations in a single mouse. B. Average T-score for each mouse represented against the number of homozygous mutations in that mouse. C. Initial T-score for each mouse plotted against the number of homozygous mutations in that mouse. D. Average learning per trial for each mouse plotted against the number of homozygous mutations in that mouse. Black horizontal lines are shown for comparison with the red best fit line.

**Supplementary figure 4.**
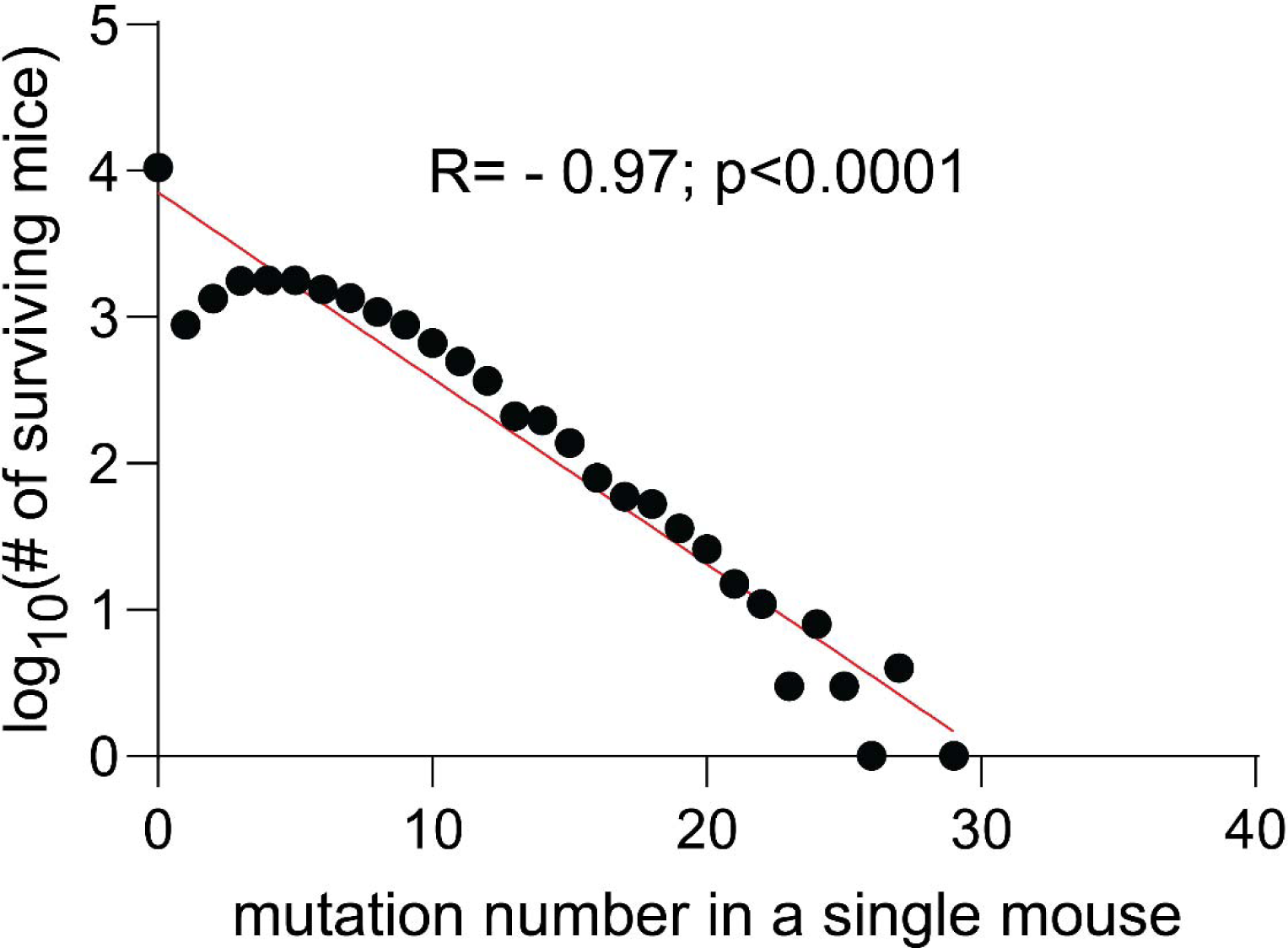
Mutation number in any single mouse in relation to the number of mice surviving with that mutation. R indicates the Pearson’s correlation coefficient.

## Notes

### Competing Interest Statement

The authors have declared no competing interest.

## REFERENCES

Adzhubei I, Jordan DM & Sunyaev SR. (2013). Predicting functional effect of human missense mutations using PolyPhen-2. Curr Protoc Hum Genet Chapter 7, Unit7 20.

Bogdanovich S, Krag TO, Barton ER, Morris LD, Whittemore LA, Ahima RS & Khurana TS. (2002). Functional improvement of dystrophic muscle by myostatin blockade. Nature 420, 418–421.

Buitrago MM, Schulz JB, Dichgans J & Luft AR. (2004). Short and long-term motor skill learning in an accelerated rotarod training paradigm. Neurobiology of learning and memory 81, 211–216.

Chebib J, Jackson BC, Lopez-Cortegano E, Tautz D & Keightley PD. (2021). Inbred lab mice are not isogenic: genetic variation within inbred strains used to infer the mutation rate per nucleotide site. Heredity (Edinb) 126, 107–116.

Cinà DP, Ketela T, Brown KR, Chandrashekhar M, Mero P, Li C, Onay T, Fu Y, Han Z & Saleem M. (2019). Forward genetic screen in human podocytes identifies diphthamide biosynthesis genes as regulators of adhesion. American Journal of Physiology-Renal Physiology 317, F1593–F1604.

Collins BE, Sweatt JD & Greer CB. (2019). Broad domains of histone 3 lysine 4 trimethylation are associated with transcriptional activation in CA1 neurons of the hippocampus during memory formation. Neurobiol Learn Mem 161, 149–157.

Cordes SP. (2005). N-ethyl-N-nitrosourea mutagenesis: boarding the mouse mutant express. Microbiol Mol Biol Rev 69, 426–439.

Drake JW, Charlesworth B, Charlesworth D & Crow JF. (1998). Rates of spontaneous mutation. Genetics 148, 1667–1686.

Dumont BL. (2019). Significant Strain Variation in the Mutation Spectra of Inbred Laboratory Mice. Mol Biol Evol 36, 865–874.

Eyre-Walker A & Keightley PD. (2007). The distribution of fitness effects of new mutations. Nature Reviews Genetics 8, 610–618.

Funato H, Miyoshi C, Fujiyama T, Kanda T, Sato M, Wang Z, Ma J, Nakane S, Tomita J, Ikkyu A, Kakizaki M, Hotta-Hirashima N, Kanno S, Komiya H, Asano F, Honda T, Kim SJ, Harano K, Muramoto H, Yonezawa T, Mizuno S, Miyazaki S, Connor L, Kumar V, Miura I, Suzuki T, Watanabe A, Abe M, Sugiyama F, Takahashi S, Sakimura K, Hayashi Y, Liu Q, Kume K, Wakana S, Takahashi JS & Yanagisawa M. (2016). Forward-genetics analysis of sleep in randomly mutagenized mice. Nature 539, 378–383.

Guenet JL. (2005). The mouse genome. Genome Res 15, 1729–1740.

Jakkamsetti V, Balasubramaniam S, Grover N & Pascual JM. (2022). Mitochondrial disease manifestations in relation to transcriptome location and function. Mol Genet Metab 135, 82–92.

Jakkamsetti V, Scudder W, Kathote G, Ma Q, Angulo G, Dobariya A, Rosenberg RN, Beutler B & Pascual JM. (2021). Quantification of early learning and movement sub-structure predictive of motor performance. Sci Rep 11, 14405.

Kanoh Y, Matsumoto S, Fukatsu R, Kakusho N, Kono N, Renard-Guillet C, Masuda K, Iida K, Nagasawa K, Shirahige K & Masai H. (2015). Rif1 binds to G quadruplexes and suppresses replication over long distances. Nat Struct Mol Biol 22, 889–897.

Kong A, Frigge ML, Masson G, Besenbacher S, Sulem P, Magnusson G, Gudjonsson SA, Sigurdsson A, Jonasdottir A, Jonasdottir A, Wong WS, Sigurdsson G, Walters GB, Steinberg S, Helgason H, Thorleifsson G, Gudbjartsson DF, Helgason A, Magnusson OT, Thorsteinsdottir U & Stefansson K. (2012). Rate of de novo mutations and the importance of father’s age to disease risk. Nature 488, 471–475.

Kuleshov MV, Jones MR, Rouillard AD, Fernandez NF, Duan Q, Wang Z, Koplev S, Jenkins SL, Jagodnik KM & Lachmann A. (2016). Enrichr: a comprehensive gene set enrichment analysis web server 2016 update. Nucleic acids research 44, W90–W97.

Kumar V, Kim K, Joseph C, Kourrich S, Yoo SH, Huang HC, Vitaterna MH, de Villena FP, Churchill G, Bonci A & Takahashi JS. (2013). C57BL/6N mutation in cytoplasmic FMRP interacting protein 2 regulates cocaine response. Science 342, 1508–1512.

Kumar V, Kim K, Joseph C, Thomas LC, Hong H & Takahashi JS. (2011). Second-generation high-throughput forward genetic screen in mice to isolate subtle behavioral mutants. Proceedings of the National Academy of Sciences 108, 15557–15564.

Lein ES, Hawrylycz MJ, Ao N, Ayres M, Bensinger A, Bernard A, Boe AF, Boguski MS, Brockway KS & Byrnes EJ. (2007). Genome-wide atlas of gene expression in the adult mouse brain. Nature 445, 168–176.

Levin SN, Riley CS, Dhand A, White CC, Venkatesh S, Boehm B, Nassif C, Socia L, Onomichi K & Leavitt VM. (2020). Association of social network structure and physical function in patients with multiple sclerosis. Neurology 95, e1565–e1574.

Long KK, O’Shea KM, Khairallah RJ, Howell K, Paushkin S, Chen KS, Cote SM, Webster MT, Stains JP, Treece E, Buckler A & Donovan A. (2019). Specific inhibition of myostatin activation is beneficial in mouse models of SMA therapy. Hum Mol Genet 28, 1076–1089.

Lucas D, Delgado-Garcia JM, Escudero B, Albo C, Aza A, Acin-Perez R, Torres Y, Moreno P, Enriquez JA, Samper E, Blanco L, Fairen A, Bernad A & Gruart A. (2013). Increased learning and brain long-term potentiation in aged mice lacking DNA polymerase mu. PLoS One 8, e53243.

Madabhushi R, Gao F, Pfenning AR, Pan L, Yamakawa S, Seo J, Rueda R, Phan TX, Yamakawa H, Pao PC, Stott RT, Gjoneska E, Nott A, Cho S, Kellis M & Tsai LH. (2015). Activity-Induced DNA Breaks Govern the Expression of Neuronal Early-Response Genes. Cell 161, 1592–1605.

Noble D, Denyer JC, Brown HF & DiFrancesco D. (1992). Reciprocal role of the inward currents ib, Na and i(f) in controlling and stabilizing pacemaker frequency of rabbit sino-atrial node cells. Proc Biol Sci 250, 199–207.

Piochon C, Kloth AD, Grasselli G, Titley HK, Nakayama H, Hashimoto K, Wan V, Simmons DH, Eissa T, Nakatani J, Cherskov A, Miyazaki T, Watanabe M, Takumi T, Kano M, Wang SS & Hansel C. (2014). Cerebellar plasticity and motor learning deficits in a copy-number variation mouse model of autism. Nat Commun 5, 5586.

Richard P & Manley JL. (2017). R Loops and Links to Human Disease. J Mol Biol 429, 3168–3180.

Schuelke M, Wagner KR, Stolz LE, Hubner C, Riebel T, Komen W, Braun T, Tobin JF & Lee SJ. (2004). Myostatin mutation associated with gross muscle hypertrophy in a child. N Engl J Med 350, 2682–2688.

Scully R, Panday A, Elango R & Willis NA. (2019). DNA double-strand break repair-pathway choice in somatic mammalian cells. Nat Rev Mol Cell Biol.

Shen EY, Jiang Y, Javidfar B, Kassim B, Loh YE, Ma Q, Mitchell AC, Pothula V, Stewart AF, Ernst P, Yao WD, Martin G, Shen L, Jakovcevski M & Akbarian S. (2016). Neuronal Deletion of Kmt2a/Mll1 Histone Methyltransferase in Ventral Striatum is Associated with Defective Spike-Timing-Dependent Striatal Synaptic Plasticity, Altered Response to Dopaminergic Drugs, and Increased Anxiety. Neuropsychopharmacology 41, 3103–3113.

Sheppard K, Gardin J, Sabnis GS, Peer A, Darrell M, Deats S, Geuther B, Lutz CM & Kumar V. (2022). Stride-level analysis of mouse open field behavior using deep-learning-based pose estimation. Cell Rep 38, 110231.

Shiotsuki H, Yoshimi K, Shimo Y, Funayama M, Takamatsu Y, Ikeda K, Takahashi R, Kitazawa S & Hattori N. (2010). A rotarod test for evaluation of motor skill learning. Journal of neuroscience methods 189, 180–185.

Tubbs A & Nussenzweig A. (2017). Endogenous DNA Damage as a Source of Genomic Instability in Cancer. Cell 168, 644–656.

Uchimura A, Higuchi M, Minakuchi Y, Ohno M, Toyoda A, Fujiyama A, Miura I, Wakana S, Nishino J & Yagi T. (2015). Germline mutation rates and the long-term phenotypic effects of mutation accumulation in wild-type laboratory mice and mutator mice. Genome Res 25, 1125–1134.

Vergouts M, Marinangeli C, Ingelbrecht C, Genard G, Schakman O, Sternotte A, Calas AG & Hermans E. (2015). Early ALS-type gait abnormalities in AMP-dependent protein kinase-deficient mice suggest a role for this metabolic sensor in early stages of the disease. Metab Brain Dis 30, 1369–1377.

Wang T, Bu CH, Hildebrand S, Jia G, Siggs OM, Lyon S, Pratt D, Scott L, Russell J, Ludwig S, Murray AR, Moresco EMY & Beutler B. (2018). Probability of phenotypically detectable protein damage by ENU-induced mutations in the Mutagenetix database. Nat Commun 9, 441.

Wang T, Zhan X, Bu CH, Lyon S, Pratt D, Hildebrand S, Choi JH, Zhang Z, Zeng M, Wang KW, Turer E, Chen Z, Zhang D, Yue T, Wang Y, Shi H, Wang J, Sun L, SoRelle J, McAlpine W, Hutchins N, Zhan X, Fina M, Gobert R, Quan J, Kreutzer M, Arnett S, Hawkins K, Leach A, Tate C, Daniel C, Reyna C, Prince L, Davis S, Purrington J, Bearden R, Weatherly J, White D, Russell J, Sun Q, Tang M, Li X, Scott L, Moresco EM, McInerney GM, Karlsson Hedestam GB, Xie Y & Beutler B. (2015). Real-time resolution of point mutations that cause phenovariance in mice. Proc Natl Acad Sci U S A 112, E440–449.

Whitlock JR, Heynen AJ, Shuler MG & Bear MF. (2006). Learning induces long-term potentiation in the hippocampus. Science 313, 1093–1097.

World Health Organization. (2006). Neurological disorders: public health challenges. World Health Organization, Geneva.

Xu D, Lyon S, Bu CH, Hildebrand S, Choi JH, Zhong X, Liu A, Turer EE, Zhang Z, Russell J, Ludwig S, Mahrt E, Nair-Gill E, Shi H, Wang Y, Zhang D, Yue T, Wang KW, SoRelle JA, Su L, Misawa T, McAlpine W, Sun L, Wang J, Zhan X, Choi M, Farokhnia R, Sakla A, Schneider S, Coco H, Coolbaugh G, Hayse B, Mazal S, Medler D, Nguyen B, Rodriguez E, Wadley A, Tang M, Li X, Anderton P, Keller K, Press A, Scott L, Quan J, Cooper S, Collie T, Qin B, Cardin J, Simpson R, Tadesse M, Sun Q, Wise CA, Rios JJ, Moresco EMY & Beutler B. (2021). Thousands of induced germline mutations affecting immune cells identified by automated meiotic mapping coupled with machine learning. Proc Natl Acad Sci U S A 118.

